# Relationships of crystallinity and reaction rates for enzymatic degradation of poly (ethylene terephthalate), PET

**DOI:** 10.1101/2023.11.05.564366

**Authors:** Sune W. Schubert, Thore B. Thomsen, Kristine S. Clausen, Anders Malmendal, Cameron J. Hunt, Kim Borch, Kenneth Jensen, Jesper Brask, Anne S. Meyer, Peter Westh

## Abstract

Biocatalytic degradation of plastic waste is anticipated to play an important role in future recycling systems. However, enzymatic degradation of crystalline poly (ethylene terephthalate) (PET) remains consistently poor. Herein, we employed functional assays to elucidate the molecular underpinnings of this limitation. This included utilizing complementary activity assays to monitor the degradation of PET disks with varying crystallinity (*X*_*C*_), as well as kinetic parameters for soluble PET fragments. The results indicate that a proficient PET-hydrolase, LCC_ICCG_, operates through an endolytic mode of action, and that its activity is limited by conformational constraints in the PET polymer. Such constraints become more pronounced at high *X*_C_ values, and this limits the density of productive sites on the PET surface. Endolytic chain-scissions are the dominant reaction type in the initial stage, and this means that little or no soluble organic product occurs here. However, endolytic cuts gradually and locally promote chain mobility and hence the density of attack sites on the surface. This leads to an upward concave progress curve; a behavior sometimes termed lag-phase kinetics.

## Introduction

The last decades have brought substantial progress in identifying and engineering enzymes capable of degrading or modifying plastics, including the widely used polyester, poly(ethylene terephthalate) (PET).^[1,2]^ Notable advancements include the discovery of a bacterium that may utilize PET waste as its sole carbon and energy source^[3]^ and the discovery and engineering of enzymes with sufficient activity for use in industrial PET-monomer recovery.^[4–8]^ However, PET-hydrolases consistently showcase high sensitivity towards the degree of crystallinity (*X*_*C*_) of the substrate.^[9–12]^

Crystalline PET comprises regions of highly ordered and densely packed PET chains, which stabilize the polymeric structure and renders it less susceptible to enzymatic degradation.^[13]^ Many post-consumer products exhibit high *X*_C_ values as a result of manufacturing processes and usage conditions.^[6,14,15]^ This poses a significant challenge for bioprocessing of PET waste. Currently, the most viable strategy to overcome the recalcitrance of crystalline PET involves implementing a pretreatment step that includes micronization and decrystallization of the PET feedstock. However, this approach involves high energy and water consumption.^[16–18]^ This situation has spurred interest in enzymes capable of degrading more crystalline polyesters, potentially drawing inspiration from the well-established ability of hydrolases to degrade naturally occurring crystalline polymers, such as cellulose^[19]^ and chitin.^[20]^

In this study, we investigated the characteristics of the PET-hydrolase, LCC_ICCG_, which is generally considered a promising candidate for large-scale industrial applications.^[1,4,21]^ To assess its catalytic activity, we employed two complementary assays described previously.^[22]^ This involved continuous pH-stat measurements to detect the release of protons resulting from cleavage of ester bonds and HPLC for the analysis of the profile of soluble, organic products. Combined analysis of these parameters for substrates with different X_C_, and special focus on the initial stage of the reaction, provided a better understanding of how LCC_ICCG_ attacks PET and the role of crystallinity.

## Results

### Progress curves

The time course for the degradation of PET disks with different *X*_*C*_ was monitored by two complementary methods, HPLC and pH-stat measurements. Our HPLC protocol was capable of quantifying six different product types in the aqueous phase. These were the three mono-aromatic compounds terephthalic acid (T), mono(2-hydroxyethyl) terephthalic acid (ET), and bis(2-hydroxyethyl) terephthalate (ETE). In addition, we could resolve one peak for oligomers with respectively two, three or four aromatic rings. Each of these peaks encompassed three molecules with either zero, one or two ethylene glycol (E) as end-groups. As an example, the fragments TET, ETET and ETETE, which all have two aromatic rings, co-eluted as a single peak. In the following, we will refer to soluble fragments with two or more aromatic rings as oligo ethylene glycol terephthalates (OETs). In contrast to the HPLC measurements, the pH-stat detected all hydrolytic reactions, irrespective of whether they produced a soluble, organic product or not.

Results in Fig. 1 show the release of protons (pH stat measurements, blue trace, left ordinate) and soluble products (HPLC measurements, symbols, right ordinate) during the hydrolysis of PET disks with different degrees of crystallinity, *X*_*C*_. Under the conditions used in this experiment, concentrations of tri- and tetra-aromatic OETs were either undetectable or near the detection limit (high nM range). Thus, Fig. 1 only includes the remaining four products (T, ET, ETE and di-aromats).

**Figure 1.**
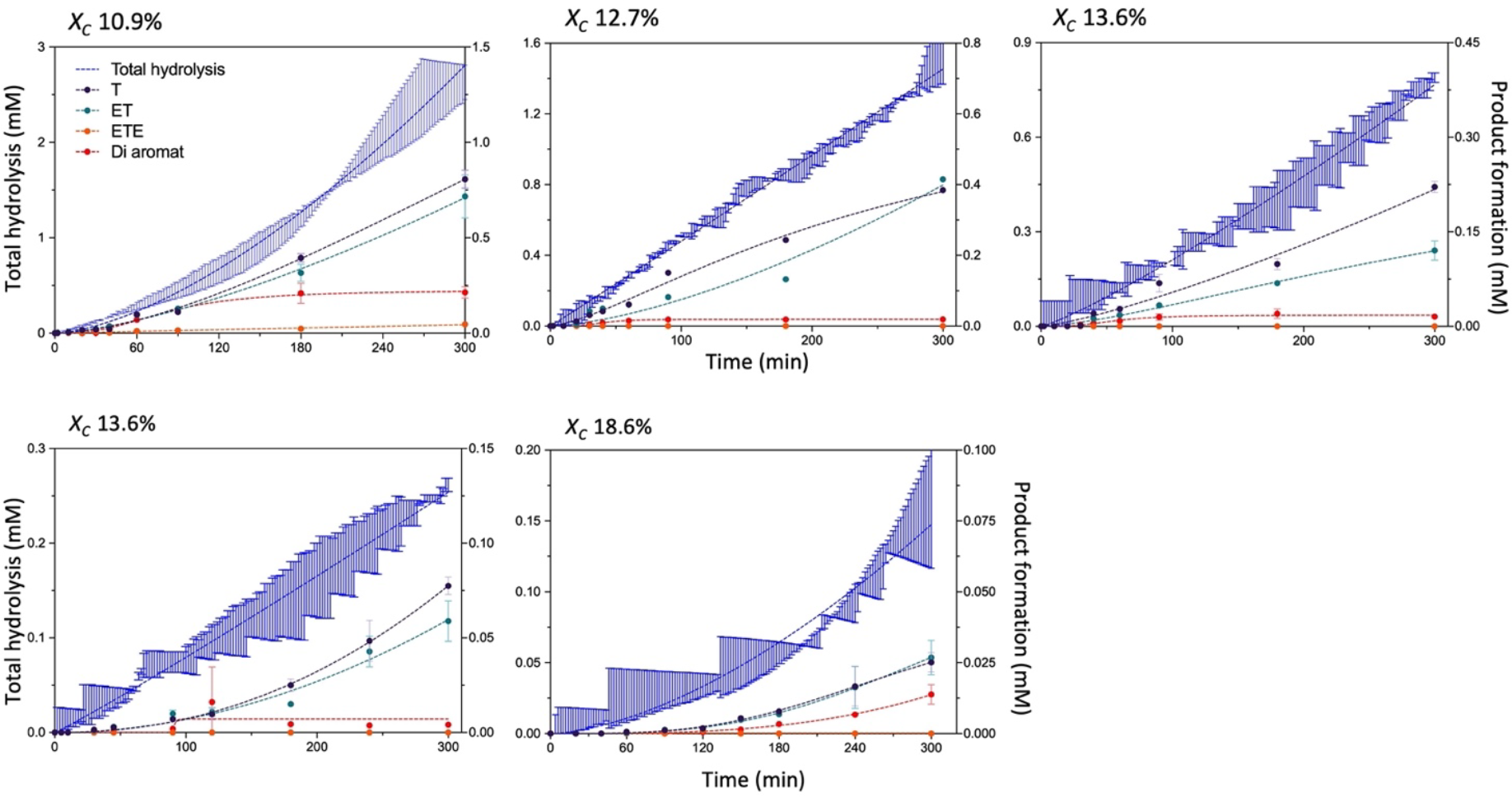
Progress curves for the activity against PET disks with varying degrees of crystallinity. Blue symbols represent the proton release (left ordinate), which provides a measure of total hydrolytic activity. Remaining curves depict the concentration of individual monomeric products and TETE, as indicated by the color code (right ordinate). Error bars were calculated as the standard deviation from two independent experiments.

To facilitate further analyses of Fig. 1, we introduced a function, R, which specified the ratio of total hydrolytic activity to soluble products. Total hydrolytic activity was defined as the number of moles of hydroxide ions delivered by the pH stat system divided by the total reaction volume. These hydroxide ions neutralized protons produced by the enzyme reaction, and we call this concentration C_H+_. The overall release of organic products, C_org_, was defined as the sum of all compounds detected by HPLC in Fig. 1, C_org_=[T]+[ET]+[ETE]+[di-aromats]. The ratio, R, may be written:

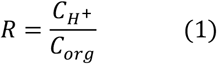

To illustrate our interpretation of this function, we note that *R*>>2 corresponds to a mode of action where the enzymatic reaction is dominated by cleavages across PET chains distant from chain ends (endo-type activity). This type of reaction generates long insoluble OETs, and hence releases protons without concurrent soluble (organic) products. Conversely, cleavage near the terminal of insoluble polymers (exo-type activity), will release commensurate amounts of protons and soluble products and hence show R ∼ 1. As a final example, complete hydrolysis of a PET chain to T, will yield *R*=2 since there are two ester bonds for each aromatic ring.

Figure 2 illustrates how *R* changes with time when LCC_ICCG_ degrades PET disks of different crystallinity. The figure also includes the functions, C_H_^+^ and C_org_, that define *R* (eq. 1). These results confirmed earlier reports on a lag phase for the release of soluble products, which became more prominent on substrates with high *X*_*C*_.^[9,23]^ For total hydrolysis, C_H_^+^, the lag phase was less pronounced and, in some cases was absent. As a result, the ratio, *R*, became larger in the initial stages of experiments for high-crystallinity substrates. Specifically, experiments with *X*_*C*_=15.6% or 18.5% yielded *R*-values around 10 after 30-90 min. Quantification of *R* earlier in the reaction was not attainable because C_org_ was too low to measure, but we clearly detected proton release, and this implies an initial reaction with high *R*-value on crystalline substrates. As argued above, this implied that initial hydrolysis of crystalline substrate was slow and dominated by an endolytic mode of action (ten or more protons were released for every soluble product). However, irrespective of the initial *X*_*C*_, *R* approached 2 after the lag phase. We emphasize that these changes in the mode of action all occur under conditions where the total degree of conversion is very low. Thus, the final degree of conversion for the measurements in Fig. 2 ranged from 0.1 to 1.7%.

**Figure 2.**
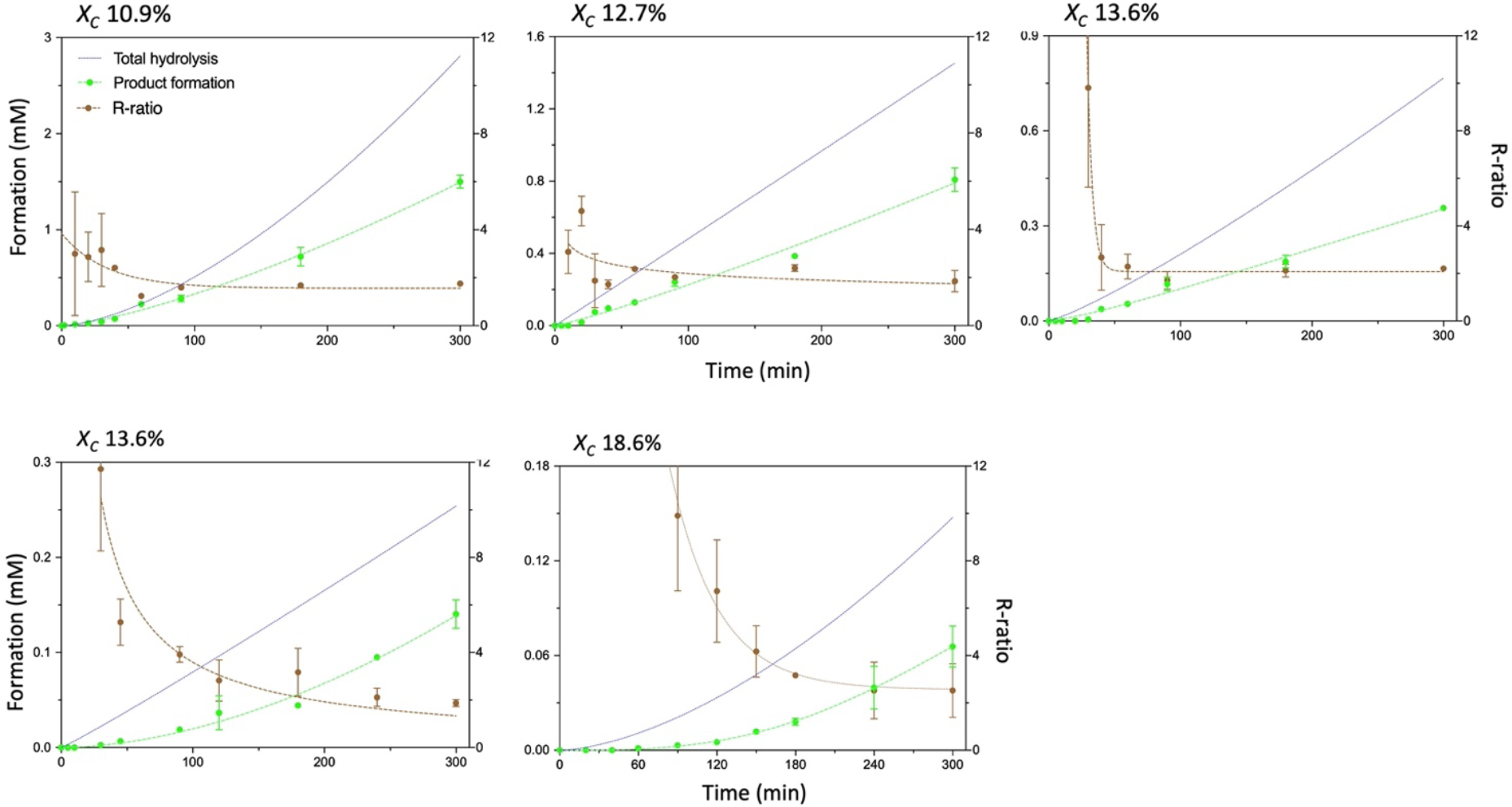
Progress curves showing the ratio R defined in eq. (1) for PET disks with varying *X*_C_ values. The left ordinate displays the sum of all organic products in Fig 1, C_org_, (green symbols) while the R-ratio (brown symbols) is displayed on the right ordinate. The blue traces replicate the proton release from Fig. 1, and are included for comparison. Elevated *R*-values in the early stages of the reaction for PET disks with higher *X*_C_ indicate that 10 or more protons are released for every soluble organic product under these conditions. Error bars represent the standard deviation from two independent experiments.

### Product profile

The results in Fig. 2 sparked interest in the initial product profile and to investigate this further, we designed a shorter time spanned experiment with a higher sampling rate. The aim of this experiment was to elucidate the profile of OETs released directly from the insoluble polymer. Thus, we used a lower enzyme concentration and higher substrate loads compared to the experiments shown in Figs. 1-2 (see Methods) to limit secondary hydrolysis of OETs in the aqueous phase. Unfortunately, the time resolution of the pH-stat system was inadequate for the desired sampling rate, and we therefore relied solely on HPLC detection in these experiments. Results in Fig. 3 show that we were able to monitor progress curves for all six product types that were detectable in HPLC (see section above). These six species were present in comparable amounts after 60 s contact time (Fig. 3A), while products with four aromatic rings dominated during earlier stages of the reaction (Fig. 3B). In accordance with Fig. 1, mono- and di-aromatic products became by far the most prevalent after several minutes of reaction, probably due to proficient secondary hydrolysis of soluble OETs in the aqueous phase (see Tab 1).

**Tabel 1.**
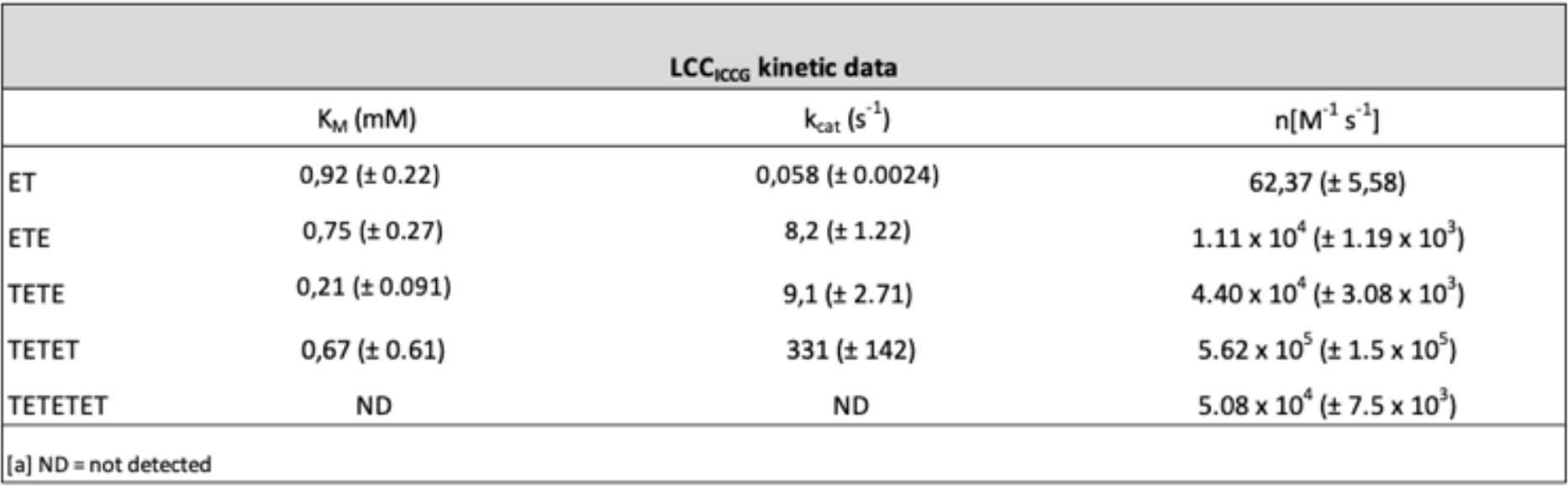
Kinetic parameters for LCC_ICCG_ acting on soluble PET fragments. Maximal turnover (k_cat_) and the Michaelis constant (K_M_) were derived from initial rate measurements (see Supplementary Material), and used to calculate the specificity constant, η=k_cat_/K_M_. Low solubility of tetra-aromatic TETETET precluded saturation for this substrate, and k_cat_ and K_M_ values could not be singled out.

**Figure 3.**
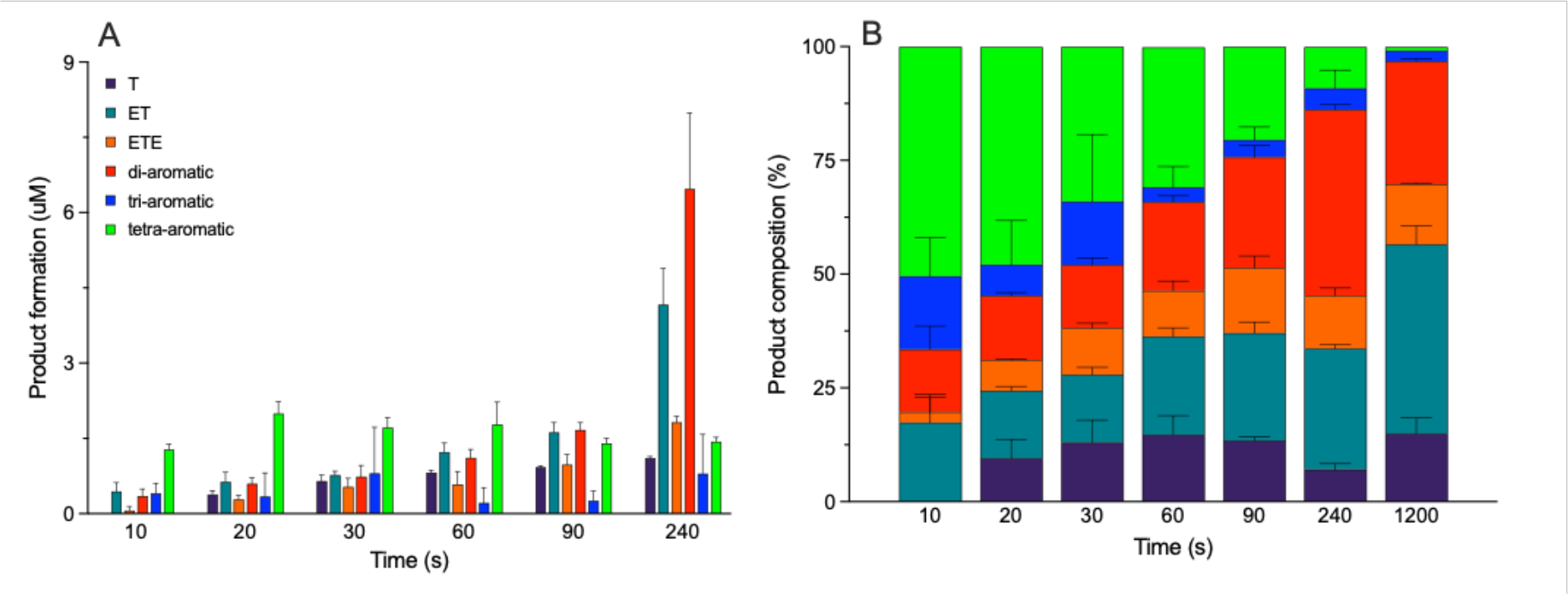
Investigation of the product profile during the early stage of an enzymatic PET hydrolysis course. Experiments were conducted at 65 °C using an enzyme load of 30 nM LCC_ICCG_ and 30 thin, amorphous PET disks (*X*_C_ = 4.6%, total mass 7 mg). These conditions were chosen to minimize the rate of secondary hydrolysis of fragments in the aqueous bulk. Panel A displays the concentrations of the six aromatic product species that could be resolved by HPLC at different time points. Panel B shows the same data illustrated as the relative abundance of each product species. Error bars represent the standard deviation from two independent experiments.

While the relative abundance of tetrameric products decreased rapidly (see Fig. 3B), their absolute value remained practically constant (1-2 µM) throughout these experiments (see Fig. 3A). This steady concentration might be attributed to low solubility and/or the rate of secondary hydrolysis in the aqueous phase. To assess these factors, we estimated the solubility of the tetrameric PET fragment under the experimental conditions used and approximated it to be 6-8 µM (see supplementary information, Fig. S1). We also determined the kinetic parameters, in terms of Michaelis-Menten kinetics, for LCC_ICCG_ acting on soluble PET fragments (Tab. 1). These results showed that the specificity constant k_cat_/K_M_ for the tetra aromatic OET was comparable to TETE but an order of magnitude lower compared to TETET. Taken together, these observations suggest that the steady concentration of tetra-aromatic product in Fig. 3B could rely on both low solubility (the aqueous concentration approaches saturation) and secondary turnover in the bulk (which is slower compared to smaller OETs). Fig. 4 illustrates a proposed reaction pathway based on the findings of the initial product profile.

**Figure 4.**
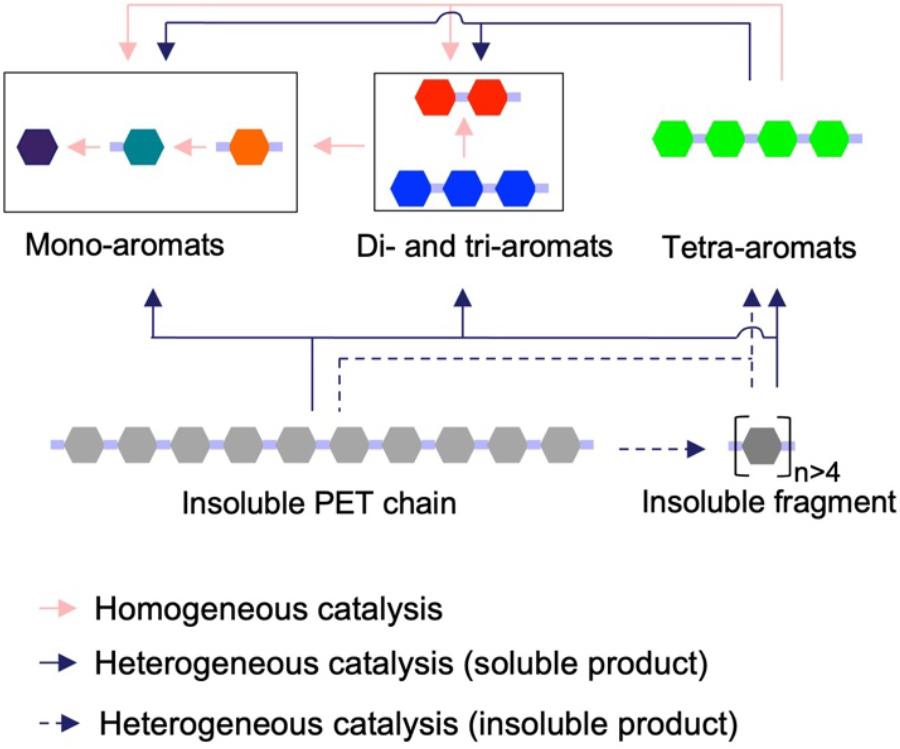
Illustration of the proposed reaction pathway for the enzymatic degradation of PET, as inferred from the product profile observed during the early stages of hydrolysis (see Fig. 3). Initially, insoluble PET chains undergo a reduction in DP due to endo-type chain scissions. This results in the formation of insoluble PET fragments. These fragments then predominantly release soluble OETs. This includes tetrameric fragments, which exhibit a low solubility (in the low μM range). Subsequently, homogeneous catalysis in the aqueous phase cleaves OETs, ETE, and ET to yield terminal T.

### Reduction in the molar mass of PET chains

In an attempt to assess the frequency of respectively endo- and exo-lytic activity we analyzed the average chain-length of PET, as a function of the degree of substrate conversion. We used a previously established approach,^[10]^ based on partial hydrolysis of a PET sample, dissolution of remaining solids in an organic solvent and detection of end-group concentration by NMR (see methods for details). Like most other methodologies, this approach suffers from the limitation that the bulk of the PET material (which is not modified by the enzyme) may dominate when all solid residue is dissolved. This has previously been discussed by Kawai et al.^[24]^ To minimize this effect, we extended the enzymatic hydrolysis to high degrees of conversion (>70% weight loss). Nevertheless, results in Fig. 5 showed an essentially constant average chain length with a degree of polymerization (DP) ranging from 225 to 250 throughout the experiment.

**Figure 5.**
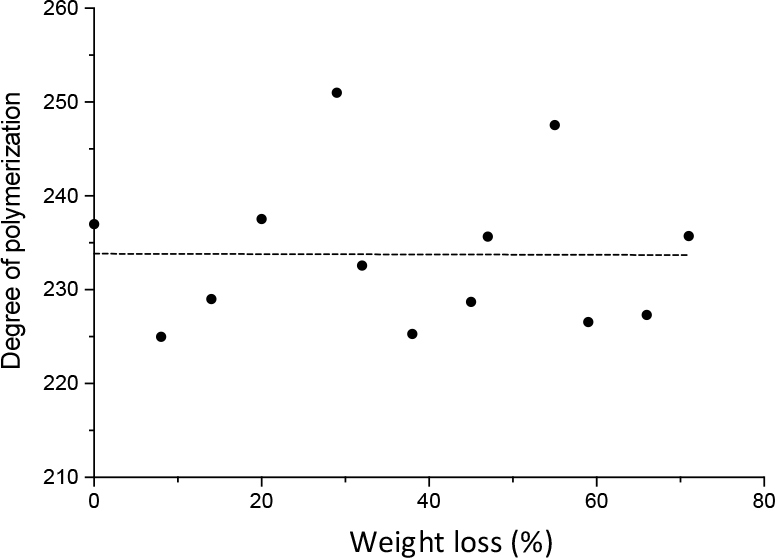
NMR analysis of polymer chain length (degree of polymerization, DP) as a function of enzymatic conversion (in % the weight loss).

## Discussion

Recent advancements in the discovery and engineering of PET-hydrolases have laid the groundwork for industrial-scale monomer recovery from polyester waste through bioprocessing^[1,21,25,26]^. Nevertheless, challenges remain for this technology to become an integral part of a circular economy. A primary concern is the limited ability of known PET-hydrolases to convert substrate with the degree crystallinity (*X*_*C*_>∼20%)^[10,27–30]^ that is typical for PET waste. While there is ample empirical evidence of the recalcitrance of crystalline PET, its molecular underpinnings are yet to be fully elucidated. Commonly proposed explanations include inaccessibility of scissile bonds, which is linked to low mobility of PET chains within or near crystalline regions.^[13,31,32]^ This interpretation is consistent with a study on the material properties of PET, which revealed suppressed chain dynamics, when amorphous samples annealed above the glass transition temperature slowly crystallized.^[33]^ Moreover, enzymatic activity has been associated with polymer chain conformation, underscoring the significance of mobility for the formation of productive enzyme-ligand complexes. ^[34,35]^

The structural data discussed above have offered valuable insights into the causes of recalcitrance, and the current work aims to investigate this topic further from a functional perspective. Our starting point was numerous observations of an initial lag phase for PET-hydrolases,^[9,22,36–38]^ which has been reported to be especially pronounced for crystalline substrates. This was also evident in the current study, as progress curves for soluble products formation (C_org,_ Fig. 2) displayed initial lag phases, which became more pronounced as the *X*_*C*_ increased. Progress curves for total hydrolysis (C_H_^+^ in Fig. 2) showcased the well-known decline in activity with increasing *X*_*C*_ (Fig. 2), but unlike for C_org_, lag-phases in C_H_^+^ were short-lived or absent. This implied that initial hydrolysis of crystalline substrates by LCC_ICCG_ was dominated by endolytic activity with limited release of organic products. Specifically, we found *R*-values above 10 for the most crystalline substrates implying that over ten protons were released initially for every soluble product.

This observation is in line with a recent study of the lag phase^[23]^ for a number of PET-hydrolases, including efficient enzymes such as LCC_ICCG_,^[4]^ PHL7,^[5]^ and duraPETase.^[7]^ The lag phases reported in this study extended up to several days for substrates with high crystallinity (>20 % *X*_*C*_). Nevertheless, soluble products emerged after even longer contact times, and in some cases, the post-lag reaction rate reached a level that was comparable (within a factor of 2-3) to the initial rate measured on the amorphous substrates (*X*_*C*_ = 12.1%). These observations and the current results collectively support the interpretation that the initial enzyme reaction involves endo-type chain scissions, which are limited by the conformation of the PET polymer. Hence, reaction only occurs if the chain is able to attain a structure that is compatible with the active site of the enzyme. Loci on the PET surface that permits this serve as “attack sites” for the enzyme but are likely to be scarcely distributed on the surface of crystalline PET due to conformational constraints, for example a fixed, linear all-*trans* crystalline structure.^[39]^

This interpretation could explain both the low rate of proton release (as attack sites are scarce) and the absence of soluble, organic products (as randomly placed attack sites across PET-chains will produce mostly long and insoluble PET fragments) (Fig. 2). A comparable explanation has been proposed by Tarazona et al.^[40]^. More importantly, this interpretation based on chain mobility and attack sites could explain the basic origin of lag phase kinetics, because breakage of one ester bond would promote local chain mobility and hence create new attack sites for the enzyme). In other words, the initial enzymatic modification gradually renders crystalline PET more reactive, and this leads to upward curvature of the progress curve (for a schematic illustration, see Fig. 6).

**Figure 6.**
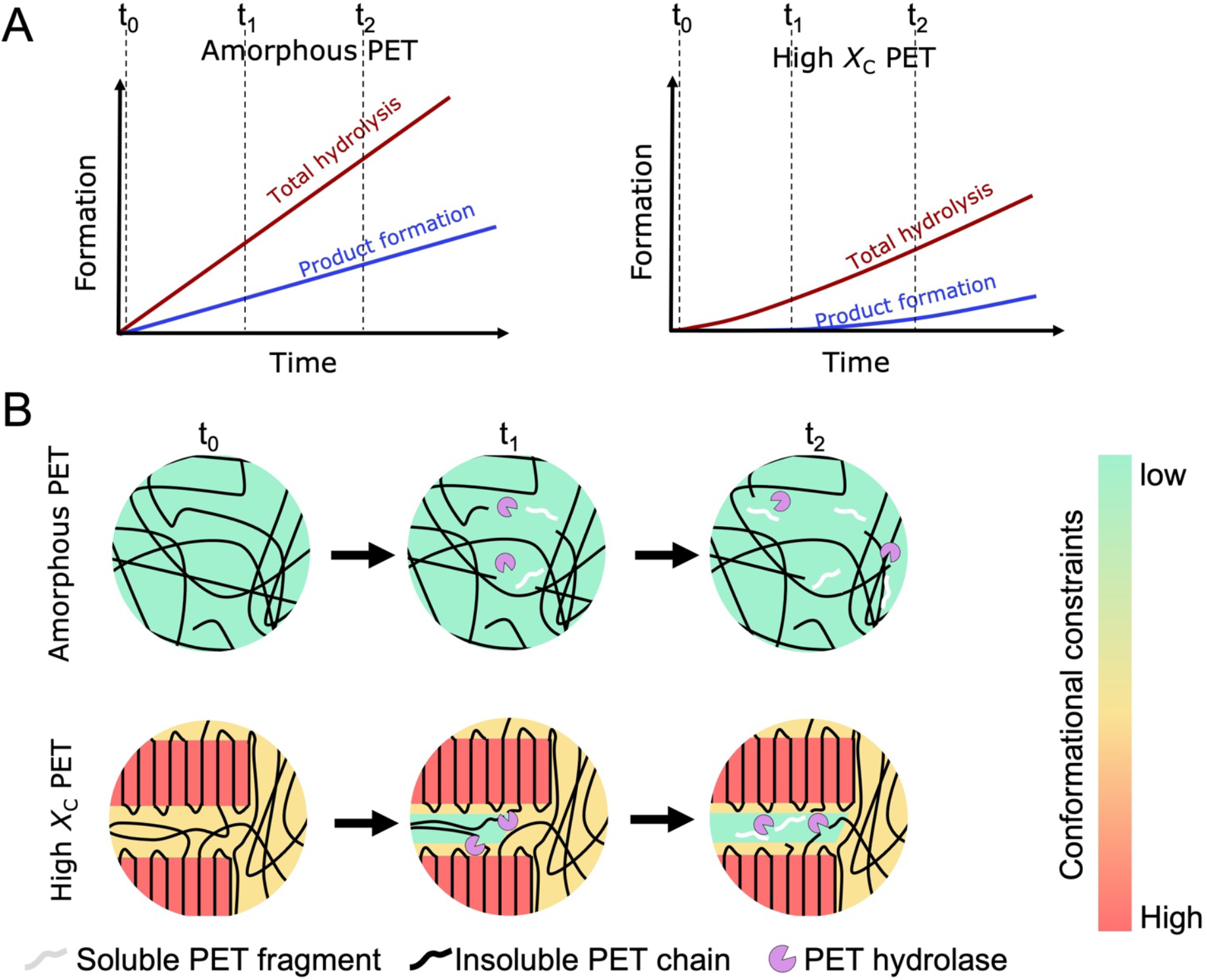
Schematic illustration of the interpretation put forward here. A) Illustrates the experimentally observed progression in total hydrolysis and product formation for PET disks with varying crystallinity. Panel B illustrates the conformational constraints that characterize PET chains in or near crystalline regions (panel B, left) and that limits enzymatic turnover. These constraints are partially alleviated by a slow, endolytic activity of LCCICCG (lower panel B, right), and this leads to a gradually increasing reaction rate, so-called lag phase kinetics. The higher mobility of chains in amorphous PET (upper panel B) underpins a faster catalytic turnover with little or no lag-phase.

The initial reaction pathway was further investigated by the short-term product-profiles shown in Fig. 3. These experiments were designed to have minimal influence of secondary OET hydrolysis in the aqueous bulk, and the results contradict the often-stated conclusion that PET-hydrolases primarily release the mono-aromatic compounds ET and ETE from chain ends of insoluble PET chains via a combined endo- and exolytic mode of action. ^[10,24,40–43]^ Thus, after 10 s, the summed concentrations of monomeric products accounted for less than 20% of soluble products. Later in the reaction, ET indeed became much more prevalent, but based on the amount of OETs released continuously (Fig. 3A), and their high turnover rates (Tab. 1), we propose that ET predominantly accumulates as the result of secondary hydrolysis of OETs in the aqueous phase. This interpretation parallels conclusions based on kinetic modeling of the degradation of nanoPET.^[22]^ Such a mode of action that predominantly proceeds via endo-type activity is also in line with the occurrence of longer soluble fragments (OETs), as observed in Fig. 3. In the simplest interpretation, where attack sites are completely randomly distributed, a random-endo pathway would generate equal amounts of all OETs and mono aromats. We did indeed observe quite comparable concentrations after e.g., 60 s, but there was a dominance of tetra-aromatic compounds in the earliest part of the experiment (∼50%) (see Fig. 3B). Interestingly, Eberl et al.^[44]^ also reported a high abundance of tetra-aromatic products in the insoluble fraction of enzyme treated PET samples. If indeed so, preference for the direct release of tetra-aromatic products could rely on either enzyme structure or chain conformation. Regarding the former, we are not aware of structural features of LCC_ICCG_ that could promote binding of tetra- aromatic moieties in the product site, and hence favor the release of such products. Alternatively, the distribution in Fig. 3 could rely on local structure and mobility of the polymer. Specifically, if a bond located four segments away from a broken ester became prone to enzymatic attack, there would be a preference for primary release of such products. Further analyses of this problem would require more experimental work, including quantification of longer fragments in the insoluble fraction of the hydrolysate.

An endo-lytic mechanism as proposed above is expected to impart rapid chain shortening to the surface layer. Nevertheless, our NMR analysis of the average chain length failed to detect any evident reduction in DP even at 70% conversion of the substrate (Fig. 5). A similar result has been reported recently,^[45]^ using another PET-hydrolase and experimental methodology. Both this and the current work used all insoluble PET residue in the analysis, and this may hamper the interpretation as it is not straightforward to assess what fraction of the residue was exposed to the enzyme and what fraction made up the (unexposed) PET bulk. However, it is notable that no changes occurred in Fig. 4, even at high degrees of conversion, and we suggest that this could be caused by the aforementioned linkage between chain mobility and attack sites. Thus, if bond breakage promotes local chain mobility, it would generate a preference for repeated attacks on the same (more flexible) strand, and it follows that this chain would be preferentially solubilized. A preference for repeated attacks on the same strand is also in accord with R∼2 (Fig. 2). Random cuts on different strands would give much higher R values because insoluble products would dominate. Further assessment of this interpretation would require data on the average chain length of insoluble PET fragments in the surface layer, and we are currently pursuing options for this type of assay.

## Conclusion

We have studied the activity of the enzyme LCC_ICCG_ on PET substrates with different crystallinity, and used the results to assess the origins of the recalcitrance of crystalline PET. A combined interpretation of current results and previous structural data suggested that conformational constraints of the PET chain, which become severe with increasing crystallinity, limit the initial enzyme reaction. This suggests that the structure of a constrained polymer rarely fits the architecture of the enzyme’s active site, and as a result, the density of productive sites (attack sites) is low on the surface of crystalline PET. This scarcity of attack sites results in a low reaction rate as seen broadly for crystalline substrates. It also explains initial dominance of endolytic reactions and insoluble products reported here (*R*≥10, Fig. 2) because activity is limited to scattered attack sites. However, slow, endolytic activity was found to change the mode of reaction even at very low degrees of conversion, and soluble products quickly became more prevalent (*R*=2, Fig. 2). We propose that this reflects a gradual and local alleviation conformational constraints. Thus, breakage of an ester bond may promote mobility of adjacent chain segments and hence the ability of these segments to combine productively with the enzyme. This produces more attack sites and leads and a type of synergy that generates an upward curvature of the progress curve. We suggest that this underpins the distinctive lag phase kinetics for crystalline substrates reported here and earlier. An interpretation based on local flexibility is also in line with the rapid decline of the *R* value (Fig. 2) for crystalline substrates because clustering of cuts would release more soluble products than random cuts, and hence drive down *R* as it is indeed observed. Finally, local alleviation of constraints could lead to a preference for repeated cuts in the same strand in accord with the observations in Fig. 5.

### Experimental Section

#### Enzymes

The PET-hydrolase LCC_ICCG_ was expressed in a heterologous system using *Escherichia coli* cells and purified according to the method described by Thomsen et al.^[46]^ The enzyme concentration was determined by measuring the absorbance at 280 nm and utilizing previously calculated molar extinction coefficients.^[47]^

#### PET substrates

PET disks with a diameter of 6 mm and weighing approximately 32 mg were generated using a generic hole punch and 1 mm thick PET sheets (Goodfellow Cambridge Ltd, Huntingdon, UK, Cat. No. ES303010). To increase the surface-to-mass ratio for PET disks subjected to NMR analysis, we incorporated thinner PET disks with a thickness of 250 µm and a weight of approximately 7 mg (Goodfellow Cambridge Ltd, Huntingdon, UK, Cat. No. ES301445).

T and ETE, which were utilized as standard samples, were purchased from Sigma. However, ET was not commercially available and was therefore produced in-house via enzymatic hydrolysis of 10 mM ETE over a period of 5 h. The purity of the ET product was determined to be > 90%, as assessed by RP-HPLC analysis.^[47]^

#### Synthesis of TETETET

A series of synthesis steps were conducted to obtain the PET fragment TETETET. This included Mono-*t*Bu terephthalate (tBu-T) which was purchased from Enamine Ltd (NJ, USA), while all other chemicals were purchased from Merck and used without further purification. Thin-layer chromatography (TLC) was performed on pre-coated silica gel on aluminum foil with a fluorescent indicator, and spots were visualized under 254 nm UV light. Flash chromatography was carried out using an automated Biotage Selekt system with pre-packed Biotage Sfär silica gel cartridges. NMR spectra were recorded at 25 °C on a Bruker Avance IIIHD 400 MHz instrument equipped with a 5 mm room temperature BBFO probe (see Fig. S2 in supplementary section for NMR spectra).

Initially, Mono-tBu terephthalate was dissolved in DCM-SOCl_2_ (9:1, 100 mL), and a catalytic amount of DMF was added. The reaction was refluxed (approximately 45 °C) under N_2_ overnight, resulting in a color change from milky white to clear light yellow. The solution was then evaporated using a rotary evaporator to yield a white solid product, which was used without further purification. Next, ethylene glycol (1 mL, 18 mmol) was dissolved in DCM-pyridine (4:1, 50 mL). Molecular sieves (4 Å) were added, and the mixture was stirred under N_2_ for 1 h. Subsequently, *t*Bu-T-Cl was added, and the solution was refluxed (45-50 °C) under N_2_ overnight. TLC analysis (EtOAc-hexane 1:4) showed the formation of *t*Bu-TET-*t*Bu (Rf 0.6) and tBu-TE (Rf 0.3). The reaction mixture was evaporated, redissolved in DCM, and extracted with water to remove excess ethylene glycol. The solution was then evaporated onto silica and dry-loaded onto a flash column. Purification was carried out with a gradient from 0 to 50% EtOAc in hexane. Following this, *t*Bu-TET-*t*Bu was dissolved in TFA-DCM (1:1, 5 mL) and stirred for 1 h at room temperature. The solution was then evaporated to dryness, yielding a white solid powder product of TET, which was used without further purification. TET was dissolved in DCM-SOCl_2_ (9:1, 20 mL), and a catalytic amount of DMF was added. The reaction was refluxed (approximately 45 °C) under N_2_ overnight, before being evaporated using a rotary evaporator to yield a white solid product, which was used without further purification. This product, tBu-TE, was dissolved in pyridine-CHCl3 (1:1, 50 mL) and 4 Å molecular sieves were added. The mixture was stirred at room temperature under N_2_ for 2 h before Cl-TET-Cl was added. The reaction proceeded over the weekend (72 h) at room temperature. TLC analysis in both EtOAc-pentane 1:1 (Rf product 0.9) and acetone-DCM 1:20 (Rf product 0.9) was conducted. The reaction mixture was filtered, evaporated, redissolved in CHCl_3_, and evaporated on silica. Purification by flash chromatography was performed using a gradient from 0-5% acetone in DCM. NMR analysis showed the target compound, albeit with a minor *t*Bu-TE impurity. Lastly, *t*Bu-TETETET-*t*Bu was deprotected in TFA-DCM (1:1, 10 mL) for 2 h at room temperature. The mixture was then carefully evaporated under 100 mbar vacuum to approximately 1 mL, followed by the addition of DCM. The solution was left at 4 °C overnight to precipitate. After decanting the supernatant, the white crystals were dried under vacuum and collected from the round-bottom flask. NMR analysis revealed the desired TETETET with trace amounts of TE (see Fig. S2 in supplementary section).

#### Isothermal crystallization

PET disks with varying substrate crystallinity were prepared through thermal annealing, and *X*_*C*_ was determined using differential scanning calorimetry (DSC) measurements in triplicates on a Pyris 1 Calorimeter (Perkin Elmer, Waltham, Massachusetts, USA). These procedures have been described in detail in a previous study.^[48]^ We note that the untreated amorphous PET disk was annealed at 85 °C for 5 min to mitigate any enthalpy relaxation caused by ageing of the polymer.

### Methods for monitoring enzymatic activity

#### Proton release detected by pH-stat analysis

The release of protons associated to cleavage of ester bonds during PET hydrolysis was monitored using the pH-stat system TitroLine® 7000 automated titrator (SI Analytics GmbH, Mainz, Germany). The reaction chamber consisted of a 10 mL vessel, equipped with a magnetic stirrer and a thermostat jacket connected to an external temperature-regulated water bath.

Activity measurements commenced with the addition of 9.5 mL of Milli-Q water to the vessel. Following this, the enzyme was added to achieve a final concentration of 150 nM, and the system was allowed to thermally equilibrate at 65 °C. The pH was then adjusted to 9.000, and the enzyme reaction was initiated by adding 9 PET disks (1 mm thickness). Subsequently, we continuously monitored the consumption of 20 mM NaOH required to maintain the pH at 9.000 over a 3-hour period. Initiation of the reaction with the substrate (rather than the enzyme) was preferred, as the enzyme stock solution influences the buffer capacity and pH of the solution. To minimize the uptake of atmospheric CO_2_ and reduce evaporation of liquid from the system, the reaction vessel was tightly sealed with a lid. To evaluate this effect and any potential leakage from the base delivery system, parallel control experiments without the substrate were conducted. Individual experiments were conducted in duplicates.

#### Product formation detected by RP-HPLC analysis

Concurrently, while performing the procedure described above for the pH-stat analysis, we retrieved subsets of the reaction mixtures for HPLC analysis. Specifically, 10 subsets containing 20 µL of the reaction mixture retrieved above the PET disks were regularly withdrawn throughout the 3-hour time course. All aliquots were diluted in 175 μL of Milli-Q water, and the reaction was quenched by the addition of 5 µL of 5 M HCl, as previously validated by Bååth et al.^[47]^

Furthermore, we investigated the product profile during the early stage of an enzymatic PET hydrolysis course. For this purpose, we deliberately exploited reaction conditions intended to minimize the catalytic rate of secondary hydrolysis of soluble products in the aqueous phase. This involved using a low enzyme dosage of 30 nM against a high substrate load consisting of 30 thin amorphous PET disks (*X*_*C*_ = 4.6%, 7 mg). Experiments were conducted at a temperature of 65 °C in a reaction volume of 2.5 mL. Over a short period and up to 10 min following the initiation of the reaction, 8 subsets of 50 µL samples were regularly withdrawn and analyzed using RP-HPLC. Individual experiments were conducted in duplicates.

Quantification and the product prodile of soluble products were analyzed by RP-HPLC using a Thermo Scientific Vanquisher system equipped with a Capital HPLC 250 mm x 4.6 mm C18 column. Samples were injected in volumes of 20 µL, employing a mobile phase consisting of 7.5 mM formic acid and 5% v/v acetonitrile at a 1:5 ratio for 7.5 min, followed by elution using acetonitrile for 12.5 min. The flow rate and column temperature were maintained at 1 mL/min and 40 °C, respectively. Analytes were detected using UV measurements at 240 nm, and peak analysis was performed using the Chromeleon Chromatography Data System software (version 7.3.1).

Mono-aromatic products (T, ET, and ETE) could be readily resolved and quantified against standard curves. Products with 2, 3, or 4 aromatic rings were also detected by HPLC; however, the different species within each of these groups (e.g., the di-aromatic products TET, TETE, and ETETE) exhibited similar retention times and eluted as one peak using the current experimental procedure. The peaks for di-, tri-, and tetra-aromatic compounds were identified and quantified using in-house synthesized standards TETE, ^[22]^ TETET, ^[22]^ and TETETET (described herein). Reactions were conducted in duplicates and included control experiments without enzyme.

#### Determination of kinetic constants

The specificity constants for LCC_ICCG_ against PET fragments ET, ETE, TETE, TETET, and TETETET was determined. Reactions were conducted in low-binding microplates (Greiner Bio-One™ 655900) in volumes of 250 uL and with substrate concentrations ranging from 0 to 2 mM. The plates were incubated in an Eppendorf thermomixer at 50°C and 1100 rpm for contact times of 10 minutes (ETE, TETE, TETET, and ETETETE) or 2 hours (ET). To prevent secondary hydrolysis, enzyme concentrations were kept low, ranging from 0.01 to 0.5 µM depending on the substrate. This approach ensured that the detected initial steady-state rate represented the hydrolysis of the respective soluble PET fragment. The specificity constant for TETETET was derived using initial reaction rates as substrate saturation was not achievable due to poor solubility (see Fig. S1). Also, to increase the solubility of the compound, we added 10% v/v DMSO to the reaction mixture, which consisted of 80 µM TETETET, 50 mM sodium phosphate buffer at pH 8, and was conducted at 50 °C and 1100 rpm. The reaction was initiated by adding LCC_ICCG_ at a final concentration of 50 nM. Quenching of the reaction was performed at specific time intervals (0, 10, 20, 30, 40, and 120 minutes) by adding 5 µL of 6 M HCl to 100 µL aliquots.

Duplicates and substrate blanks (for quantification of autohydrolysis) were included, and all reactions were quenched and analyzed by RP-HPLC as described above.

#### Assessing the solubility of PET fragments

A study was conducted to compare the solubility of various PET fragments released during enzymatic PET hydrolysis. To achieve this, concentration gradient from either 0-80 µM or 0-1 mM comprising T, ET, ETE, TET, TETE, TETET, ETETETE or TETETET PET fragments was prepared. Samples were adjusted to contain similar concentrations of DMSO at either 0.7% v/v or 1.5% v/v. These prepared PET fragment samples were introduced to a 96-well plate containing the reaction buffer used for experiments in Figs. 1 and 2. The plates were subsequently incubated at 65 °C for 1 h. After the incubation period, absorbance was measured for turbidity at 600 nm using spectrophotometric measurements in a plate reader (Molecular Devices SpectraMax Paradigm, San Jose, USA).

#### NMR analysis of partially degraded PET disks

NMR analysis was conducted on partially degraded PET disks (250 µm thickness) to study the DP as a function of the mass conversion of the PET disk. For this purpose, individual reactions were conducted using PET disks with an enzyme load of 150 nM at 65 °C in a reaction volume of 1 mL. Over a 72-hour duration, the PET disks were regularly withdrawn, dried, and subjected to gravimetric analysis to determine weight loss. Subsequently, the enzyme-treated PET disks, along with an untreated control sample, were dissolved in hexafluoroisopropanol (HFIP, ≥ 99%, Merck KGaA, Darmstadt, Germany) and stored at room temperature for 48 h to ensure complete dissolution. Aliquots of 8 µL of the dissolved PET solutions, with a final concentration of 15.0 mg/mL, were transferred into a 3 mm NMR tube and mixed with chloroform-d (CDCl_3_, 99.5%) to a final volume of 150 µL.

All 1H solution NMR measurements were carried out at 25 °C using a Bruker Avance 800 MHz spectrometer (Bruker Biospin, Germany) operating at a 1H frequency of 800.09 MHz and equipped with a 5 mm TCI cryoprobe. The 1H NMR spectra were acquired using the zg experiment with a 1H 90° pulse length of 6.8 μs and a recycle delay of 15 s. A total of 64k data points spanning a spectral width of 20 ppm were collected in 256 transients. The spectra were processed using Topspin (Bruker). An exponential line broadening of 0.3 Hz was applied to the free induction decay prior to Fourier transformation, and the baseline was corrected. Individual baseline correction was applied for the signals at 4.49, 4.00, and 3.89 ppm using Matlab (The MathWorks Inc., Natick, USA). The intensities of all signals were measured in Matlab.

Estimation of the DP was performed as previously described by Wei et al.^[10]^, which involved analyzing the peaks in the spectral regions of 7.8–8.2 ppm and 3.8–4.8 ppm. Integration ratios are provided under each signal, calculated relative to the CH_2_ signal at 4.0 ppm.

## Supporting information

Supplementary section

## Funding

This work was supported by the Novo Nordisk foundation Grant NNFSA170028392 (to P. W.). In addition, this work was supported by a grant (Project no. 40815) from Villum Fonden, The Villum Experiment Programme

## Conflict of Interest

K.B., K.J., and J.B. work for Novozymes A/S, a major manufacturer of industrial enzymes.

## Supplementary information

**Figure S1.**
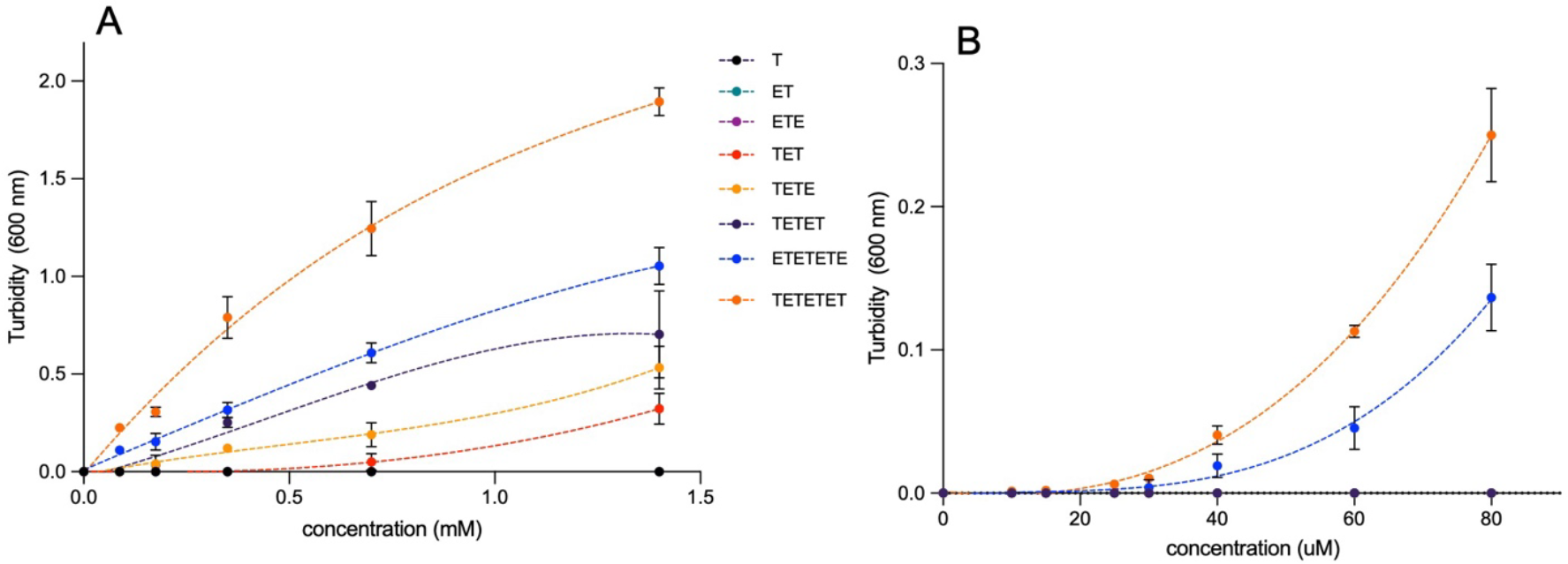
Comparison of the solubility of various PET fragments released during enzymatic PET hydrolysis. A concentration gradient comprising different species of mono-, di-, tri-, and tetrameric PET fragments was introduced to a 96-well plate containing the reaction buffer used in Figs. 1 and 2. The plates were incubated at 65 °C for 1 h, after which the turbidity was measured at 600 nm. This signal served to assess the degree of solubility of the PET fragments. As several PET fragments were stored in DMSO stock solutions, we adjusted the concentration of all samples to contain similar concentrations of DMSO. Figure A) included 0.7% v/v DMSO, while this value was 1.5% for figure B. We acknowledge that the presence of DMSO likely enhanced the solubility of the analyzed PET fragments. In scheme A showing concentrations of the PET fragments in the 0-80 nM range, signals were only detected for TETETET and ETETETE, from which their solubility levels were estimated to be roughly 4-10 µM and 25-35 µM, respectively.

**Figure S2.**
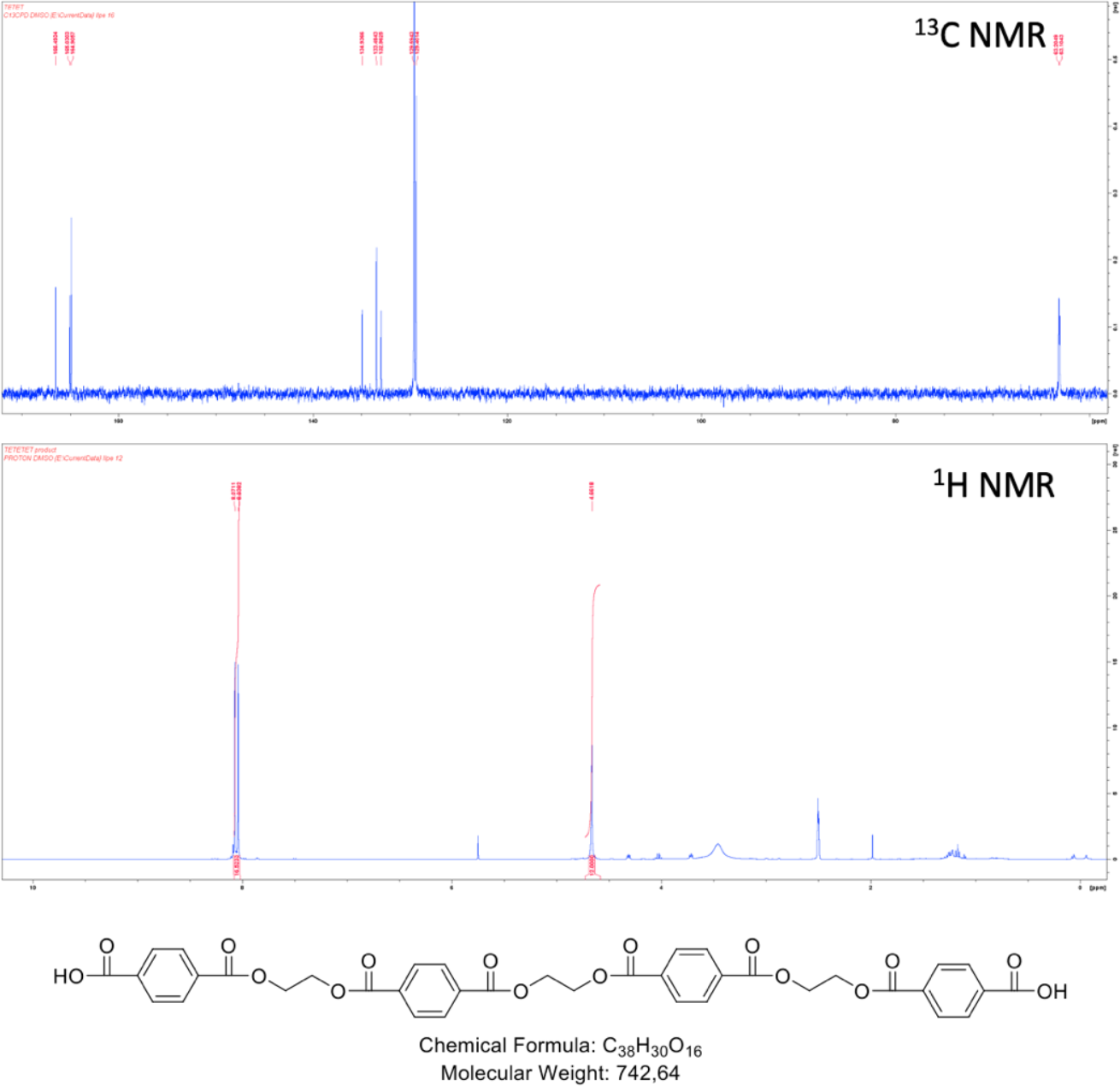
Characterization of TETETET. Upper panel show the ^13^C NMR spectra for the in-house synthesized PET fragment while the lower panel show the ^1^H NMR spektra.

## Graphical Abstract

**Graphical abstract.**
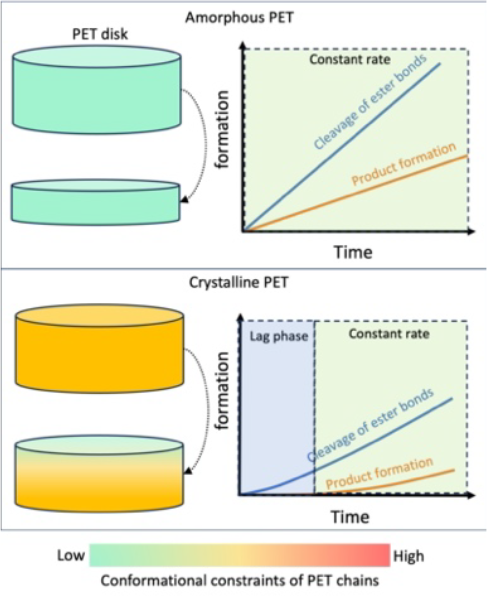
Herein, we demonstrate that LCC_ICCG_ operates via an endolytic mode of action and that its activity is limited by conformational constraints in the PET polymer. However, endo-type cuts locally promote chain mobility and hence the density of attack sites on the surface. This gradually promotes formation of soluble product

