## Supplementary section for "Relationships of crystallinity and reaction rates for enzymatic degradation of poly (ethylene terephthalate), PET"

### Supplementary information

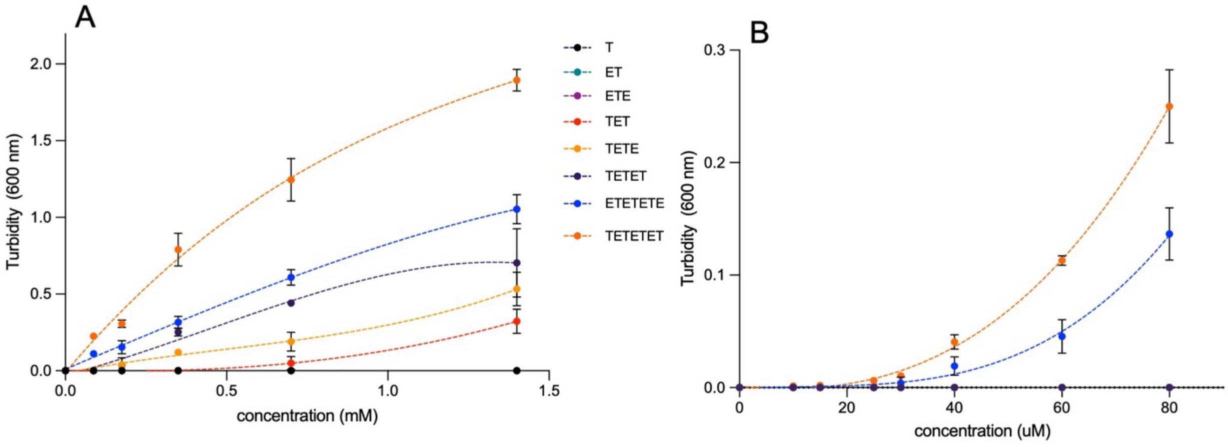


**Figure S1.** Comparison of the solubility of various PET fragments released during enzymatic PET hydrolysis. A concentration gradient comprising different species of mono-, di-, tri-, and tetrameric PET fragments was introduced to a 96-well plate containing the reaction buffer used in Figs. 1 and 2. The plates were incubated at 65 °C for 1 h, after which the turbidity was measured at 600 nm. This signal served to assess the degree of solubility of the PET fragments. As several PET fragments were stored in DMSO stock solutions, we adjusted the concentration of all samples to contain similar concentrations of DMSO. Figure A) included 0.7% v/v DMSO, while this value was 1.5% for figure B. We acknowledge that the presence of DMSO likely enhanced the solubility of the analyzed PET fragments. In scheme A showing concentrations of the PET fragments in the 0-80 nM range, signals were only detected for TETETET and ETETETE, from which their solubility levels were estimated to be roughly 4-10 μM and 25-35 μM, respectively.

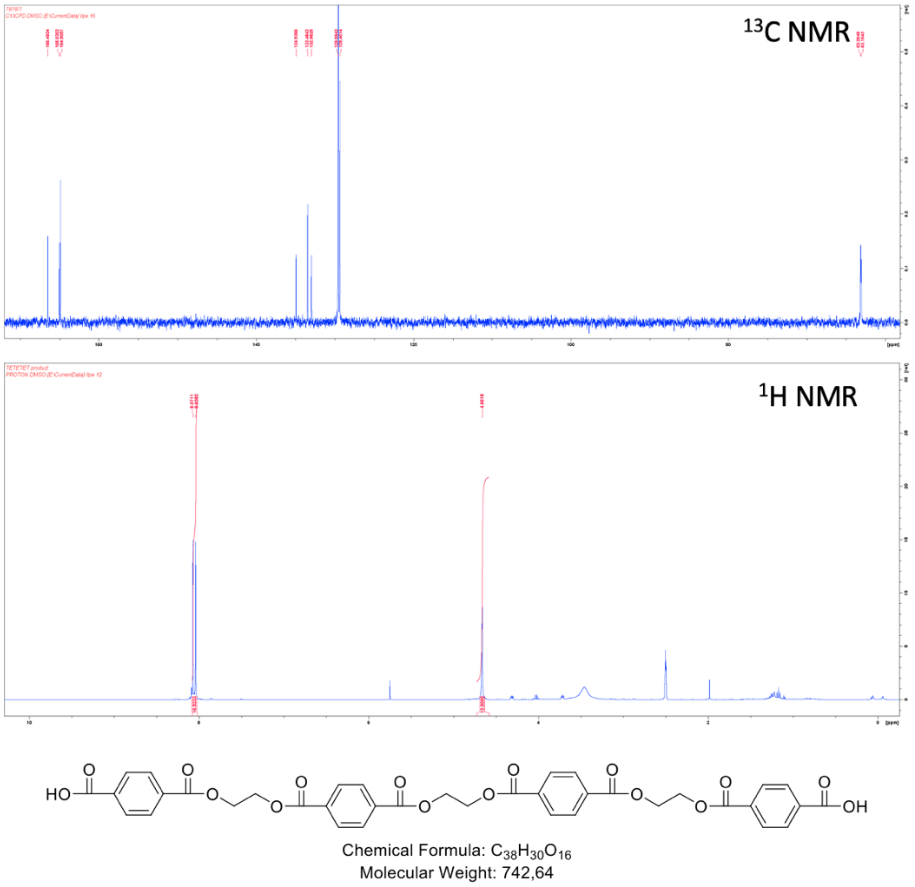


**Figure S2.** Characterization of TETETET. Upper panel show the $^{13}C$ NMR spectra for the in-house synthesized PET fragment while the lower panel show the $^{1}H$ NMR spektra.

WILEY

#### Graphical Abstract

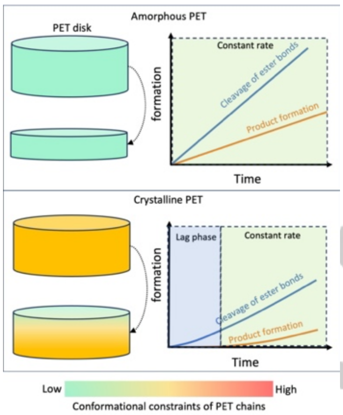


**Graphical abstract** Herein, we demonstrate that $LCC_{ICCG}$ operates via an endolytic mode of action and that its activity is limited by conformational constraints in the PET polymer. However, endo-type cuts locally promote chain mobility and hence the density of attack sites on the surface. This gradually promotes formation of soluble product

WILEY-VCH
